# In vitro pathogenicity evaluation of deep intronic variants for recessive genetic retinal diseases

**DOI:** 10.64898/2026.09.18.752412

**Authors:** Marianna Weener, Kevin Valestil, Emily Place, Sudeep Mehrotra, Daniel Navarro, Hilary Scott, Kinga Bujakowska, Eric Pierce

## Abstract

**Purpose:** Deep intronic variants (DIVs) that activate cryptic exons (CEs) cause ∼20% of inherited retinal disease (IRD) cases, yet their pathogenicity remains difficult to predict. We measured the splicing effects of rare DIVs in patients with recessive IRD who each carry one confirmed pathogenic allele.

**Methods:** We tested 640 rare DIVs from 76 patients using a high-throughput splicing assay (HTSA): a split-GFP minigene separated by an SMN1 gene intron into which 270-bp sequences flanking each DIV were cloned. The plasmid library was transfected into HEK293T cells, minigene transcripts were amplified by RT-PCR and sequenced, and intron sequences spliced between the two GFP exons were identified and quantified.

**Results:** Ninety-eight variants activated a CE more than 100-fold relative to the matched reference oligo and were classified as pathogenic in this experimental setting, with validation in a longer genomic context; 26 variants were classified as VUS. Only 6 of 90 variants (6.6%) were predicted to be pathogenic by SpliceAI. The results confirmed the diagnosis for 50 of 78 patients (64%).

**Conclusion:** Overall, 19.4% of tested DIVs caused CE activation, indicating that a substantial portion of undiagnosed patients with recessive IRD carry a pathogenic intronic variant that current prediction algorithms miss.

## Introduction

Inherited retinal diseases (IRDs) are a clinically and genetically heterogeneous group of degenerative disorders and a leading cause of irreversible blindness in working-age adults. Although more than 280 IRD genes have been described and next-generation sequencing has transformed molecular diagnosis, a substantial fraction of clinically well-characterized patients remains genetically unsolved. In the Ocular Genomics Institute (OGI) cohort and in comparable series, single-nucleotide variants and small indels account for roughly 56% of solved cases and copy-number variation for a further ∼9–10%, yet approximately 13% of families carry only a single convincing pathogenic allele in a recessive gene that matches the phenotype, and close to 20% remain fully unsolved even after whole-genome sequencing (WGS).^1–3^ When a single high-impact loss-of-function allele is found in a recessive IRD gene that fits the clinical picture, the most ‘low hanging fruit’ hypothesis is that a second, disease-causing allele lies in a region that standard analyses overlook, most often the non-coding, intronic space of the same gene.^4–6^

Deep intronic variants (DIVs) that create or strengthen splice signals can activate a cryptic exon (CE, also termed a pseudoexon): an intronic segment that becomes aberrantly included in the mature transcript, typically introducing a frameshift or premature termination codon and abolishing gene function.^7–9^ Such variants are, by construction, invisible to exome sequencing and are easily discarded by WGS filtering pipelines that prioritize coding and canonical splice-site changes. Over the past decade, individual deep intronic and pseudoexon-activating alleles have been described in a growing list of IRD genes, including *ABCA4*,^10–18^ *USH2A*,^18–21^ *CEP290*,^18,22^ RPGRIP1,^23^ TULP1,^24^ *OFD1*,^25^ *POC1B*,^26^ and *RPGR*,^27^ and in Mendelian disease more broadly *CLRN1,*^28^ *BBS,*^29^ *CFTR,*^30^ *ETFDH,*^31^ *SCN1A,*^32^ and, among others.^33–34^ The same diagnostic trajectory is unfolding across genetic medicine, from X-linked intellectual disability^31^ to Duchenne muscular dystrophy,^35–36^ hemochromatosis type 4,^37^ and chronic granulomatous disease.^38^ Collectively, these reports suggest that pseudoexon activation is a recurrent and under-recognized disease mechanism.^39–42^

The central obstacle is interpretation. A rare DIV discovered in trans with a known pathogenic allele is only a hypothesis until its effect on splicing is demonstrated. In silico splicing predictors SpliceAI, Pangolin, SQUIRLS,^41^ CI-SpliceAI, SPiP, MaxEntScan,^42^ CADD,^43^ and PDIVAS^44^ were trained predominantly on variants at or near canonical splice sites and perform poorly deep within introns, where the sequence determinants of cryptic splice-site usage are incompletely understood.^45–53^ In practice, the overwhelming majority of DIVs receive scores at or near zero, so they are filtered out before a human ever evaluates them.^54^ This creates a circular problem: because predictors flag few deep intronic variants, DIVs appear to contribute little to disease, which in turn discourages their systematic testing.^51–52^ Breaking this cycle requires an unbiased, experimental readout of splicing that does not depend on prior computational triage.

High-throughput splicing assays (HTSAs) based on minigene reporters provide this readout, measuring the splicing consequence of hundreds of candidate variants in parallel.^51^ Here we apply an HTSA built on a split-GFP *SMN1*-intron minigene adopted from Cheung R ^55^ and modified in our lab^56^ to systematically evaluate rare DIVs identified in genetically unsolved recessive IRD patients who each carry one confirmed pathogenic or likely pathogenic (P/LP) allele. By assaying 640 patient-derived DIVs across 52 IRD genes, quantifying cryptic-exon activation against matched reference sequences, and validating a representative subset in an extended genomic context and by family segregation, we ask three questions: how large a share of these “second-hit candidate” DIVs actually alter splicing, how well current predictors identify them, and how many patients can be moved from unsolved to solved. Our results indicate that deep intronic pathogenicity is considerably more common than prediction-based estimates imply, and that empiric testing is presently indispensable for its recognition.

## Materials and Methods

*Patient sequencing and annotation.* Whole genome sequencing was performed at MEE genomics core using methodology described previously.^1^ Sequencing data was aligned to human genome 38 and variants were called using Genome Analysis Toolkit (GATK) HaplotypeCaller package version 3 ^57^ (https://software.broadinstitute.org/gatk/) and annotated using VEP^58^ and VCFAnno.^59^ CNV predictions were produced using gCNV.^57^

*Intronic variants selection for cryptic exon library and oligo design*. Patients’ cases were chosen basing on clear clinical picture of IRD, inheritance pattern and enough diagnostic data. Variant search was inside the GEDI v.7 gene list with MAF ≤ 0.01 (allele frequency), All passing variants (quality reads), AR inheritance. List of variants was filtered for all 1 hits for AR IRDs which fits all the listed criteria: a) ClinVar – P/LP, VUS, conflicting interpretation, unknown b) Damaging in at least 2 out of 4 prediction algorithms c) Absence in homozygous state in GnomAD and Exac. After comparison with the clinical picture most relevant possibly causative rare deep intronic variants were chosen into the variant list.

*Deep intronic variants* (DIVs) in known IRD genes that were 50 base pairs (bp) or more from the nearest exon were selected as possible candidates for the splicing assay from the patients with already confirmed 1 pathogenic or likely pathogenic mutation in one of IRD genes in trans position. 270 bp intron oligo and 15 bp flanking adapter sequences containing AgeI and NheI restriction sites, introns included the AgeI and NheI restriction sites were excluded, we used Broad developed software, SEQR^60^ to prioritize rare deep intronic variants in these introns (MAF <0.01) found in the unsolved IRD subjects. In total 640 variants in 52 IRD genes were selected (Supplementary Table S1). Each variant was staggered 50 bp to the left and 50 bp to the right creating 3 oligos for each intronic variant for checking to exclude the nearest genomic context influencing on splicing (Figure 1B). A pool of ssDNA oligos synthesized on silicone array-based method was purchased from a commercial vendor (TWIST Biosciences, USA).

**Figure 1.**
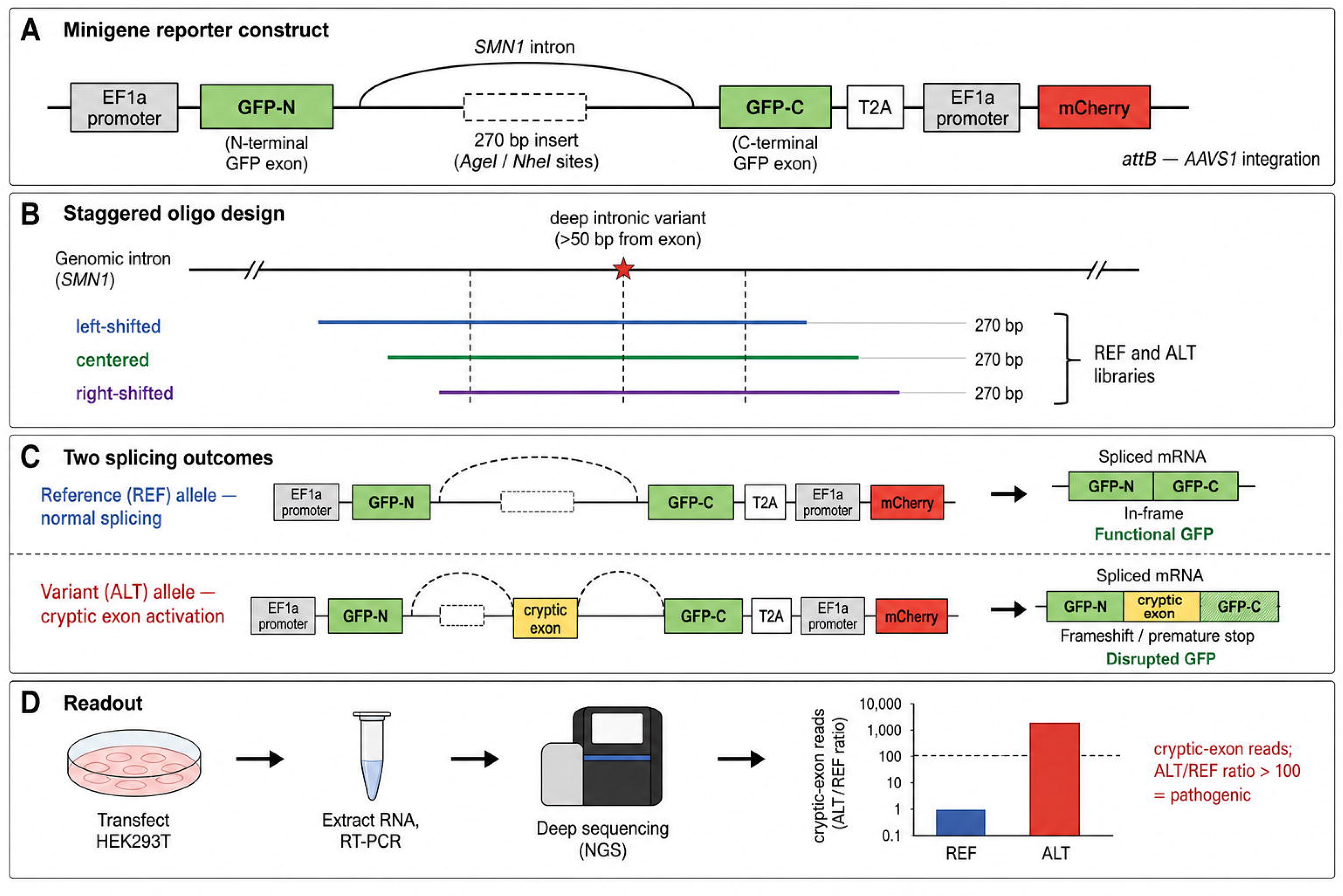
High-throughput splicing assay (HTSA) design. (**A**) Split-GFP SMN1-intron minigene reporter: correct splicing joins GFP-N to GFP-C and reconstitutes functional GFP; inclusion of a cryptic exon between the GFP halves disrupts the reading frame. (**B**) Staggered 270-bp oligonucleotide design (centered, left-shifted, and right-shifted by 50 bp) for matched reference (REF) and variant (ALT) libraries. (**C**) Splicing outcomes scored by RT-PCR and deep sequencing of transcripts spliced between the two GFP exons. (**D**) Quantification of cryptic-exon inclusion as the ALT/REF read ratio normalized to reads per million within each pool.

*High throughput splicing assay (HTSA) design*. We have adapted a published approach from Cheung and colleagues,^55^ who kindly shared with us the *SMN1* intron backbone plasmid with split GFP-T2A-mCherry, the Bxb1 integrase plasmid and the HEK2937-RCA7 cell line, that provide a chromosomal landing pad site which allows stable expression of splicing reporter library at the adeno-associated virus integration site 1 (AAVS1) safe harbor locus.^61^ In our design, we altered the *SMN1* split GFP-T2A-mCherry plasmid, where we decoupled the mCherry and GFP expression to be able to study out-of-frame exons. For the uniform expression of both fluorophores, we introduced EF1alpha promoters in front of the split GFP and mCherry genes. This altered vector we call HTSA vector. The vector also contains attB sites for integration into the AAVS1 locus of the HEK293T-RCA7 landing pad cell line, as well as a promoter-less puromycin resistance cassette. HEK293T-RCA7 contains an attP site and the promoter that allows for the site-specific recombination of the HTSA insert and coupling of the promoter and the puromycin resistance gene.

*HTSA library design and cloning*. We designed two oligo libraries: 1) control (REF) library containing 640 study intron variants with reference sequences; and 2) variant (ALT) library containing matching 640 study deep intronic variants staggered in three left, centered and right 50 bp shift, with altered sequences of study variants. This approach led to 3,840 sequences in total. Sequences that included the AgeI and NheI restriction sites were excluded. The library was amplified with Fwd-AgeI primer: 5’-ACGGCCAGTACCGGT-3’ and NheI-Rev primer 5’-GCTATGACCGCTAGC-3’ with a high-fidelity polymerase (KAPA HiFi HotStart ReadyMix, Roche) for 14 cycles and cloned to the HTSA vector by the AgeI and NheI restriction cloning.

Due to oligonucleotide length limitation of 270 bp and unknown placement of the cryptic exons this in vitro experiment caused by manufacturing technological process, we used staggering approach for both reference (REF) and altered (ALT) oligos library design to improve accuracy of detecting variants that cause splicing anomaly (Figure 1B). Maximum allowed size of oligo libraries was 300 bp, after removing 15 bp adaptors from each side, we had 270 bp oligonucleotide context, which was not enough to check the cell environment splicing machinery activity. That’s why three versions of oligos were synthesized: centered, staggered left and staggered right (Figure 1B) making it 3,840 oligos in two separate libraries (REF and ALT).

*Cell culture*. After NGS confirmation of the desired diversity (Novaseq 6000, Illumina), the libraries were transfected into the HEK293T cell line.

*Genomic DNA extraction and deep sequencing*. gDNA extraction was performed with a commercial kit (Qiagen All-Prep Kit) and the integrated fragments were PCR amplified with HTSA-NGS-F primer 5’-CTATATATAGCTATCTATGTCTACCGGT-3’ and a commercial kit (Qiagen All-Prep Kit) and the integrated fragments were PCR amplified with HTSA-NGS-R primer 5’-GGCTGGAACTCTTGCGCTAG-3’ with a high-fidelity polymerase (Primestar GXL Polymerase, Takara) for a maximum of 20 cycles. The amplicons were multiplexed and deep sequenced with a targeted depth of >500 reads per oligo (250×250 pair-end reads, Novaseq 6000, Illumina). Primers for phasing were designed using Primer 3 software (ELIXIR - European research infrastructure for biological information).

*Clinical investigations and genetic counselling* were performed at Mass Eye and Ear (MEEI) and included Visual acuity and Goldman visual fields check, IOP (intraocular pressure) measurement, ophthalmoscopy, OCT (optical coherent tomography) and FAF (fundus autofluorescence) measurement.

*Data analysis and statistics*. Deep intron sequence reads were determined by NGS and matched to the oligo sequences in the corresponding library, counted and normalized by coverage (reads per million in each pool). RT-PCR sequencing data was providing reads count for each oligonucleotide fragment. Ratio of ALT/REF was measured. Three biological replicates were done for each comparison. All figures were prepared with Microsoft Excel, Graphpad Prism 8, Biorender, and Adobe Illustrator.

## Results

### Patient cohort and candidate variant selection

From 237 patients with clinically confirmed IRDs at the OGI, we selected individuals with a well-documented phenotype and family history consistent with recessive retinal degeneration that remained unsolved after WGS. A case was eligible if exactly one P/LP variant had been identified in a recessive IRD gene matching the phenotype and no structural variant explained the disease. This yielded 78 carrying a single P/LP allele and at least one rare DIV in the same gene in trans position. For each patient we prioritized rare DIVs (minor allele frequency ≤ 1×10, consistent with Variant Curation Expert Panel guidance^62^) located at least 50 bp from the nearest annotated exon, excluding peri-exonic variants that are evaluated separately.^41^ In total, 640 deep intronic variants across 52 IRD genes were carried forward for functional analysis.

### A split-GFP minigene assay resolves cryptic-exon inclusion

We adapted a published split-GFP HTSA in which the two halves of GFP (GFP-N and GFP-C) are separated by an *SMN1* gene intron carrying a cloning slot for a 270-nucleotide test sequence (Figure 1A).^55–56^ Correct splicing joins GFP-N to GFP-C and reconstitutes functional GFP; inclusion of a cryptic exon between the GFP halves disrupts the reading frame and can be detected by RT-PCR and sequencing of the transcripts spliced between the two GFP exons. To overcome the 270-bp oligonucleotide length limit and the unknown position of the cryptic splice sites relative to each variant, every variant was represented by three staggered 270-bp oligonucleotides centered, left-shifted, and right-shifted by 50 bp (Figure 1B), so that a cryptic exon falling near an oligo boundary would still be captured in at least one context. Matched reference (REF) and variant (ALT) libraries were synthesized, giving 3,840 sequences in total, and cloned into the HTSA vector (Figure 1C). Library representation after cloning was 98% (REF) and 97% (ALT). Libraries were transfected into HEK293T cells, RNA was harvested at 48 h, and spliced products were quantified by deep sequencing, normalizing read counts to reads per million within each pool (Figure 1D). Three biological replicates were performed per comparison, and per-variant ALT/REF ratio of cryptic-exon reads was used as the pathogenicity metric.

### Cryptic-exon activation is common among candidate deep intronic variants

Applying a pre-specified classification ALT/REF ratio > 100 for pathogenic (P/LP), 2–100 for uncertain (VUS), and < 2 for benign. 124 of 640 variants (19.4%) activated a cryptic exon, comprising 98 pathogenic variants (15.3%) and 26 VUS (4.1%); the remaining 516 variants (80.6%) showed no splicing defect. The requirement for a matched REF comparison is important: because transient transfection introduces hundreds of plasmid copies per cell, low-level aberrant splicing is occasionally observed even from reference sequences, and the ALT/REF ratio controls for this background rather than relying on an absolute ALT signal alone. The 100-fold and 2-fold cut-offs were deliberately conservative; the large gap between the pathogenic and benign thresholds creates an uncertainty zone (the VUS class) rather than forcing a binary call, and the downstream validation (below) was designed to test whether variants at these boundaries behave as the thresholds predict. Read depth exceeded 500 reads per oligonucleotide, so the ratios for the great majority of variants rest on hundreds to thousands of reads per replicate.

Cryptic exons were activated across 28 of the 52 genes tested (53.8%) (Figure 2A). The burden was highly uneven: *USH2A* (29 pathogenic CEs), *EYS* (15), *ADGRV1* (9), *VPS13B* (7), *ABCA4* (5), and *CDH23* (5) together accounted for 70 of 98 pathogenic variants (71%). These are among the longest genes in the panel, with correspondingly large intronic search space, so the concentration of pathogenic DIVs in these genes reflects both their intron length and the number of candidate variants they contributed. Notably, hit rates differed markedly by gene: *EYS* yielded pathogenic CEs in only ∼5% of the 292 variants tested, whereas several genes with short introns and thus few rare suspicious deep intronic variants *in trans* with the revealed exonic variant: *ACO2*, *BBS1*, *FLVCR1*, and *NR2E3* activated cryptic exons in nearly every variant assayed, indicating that per-gene splice susceptibility varies substantially.

**Figure 2.**
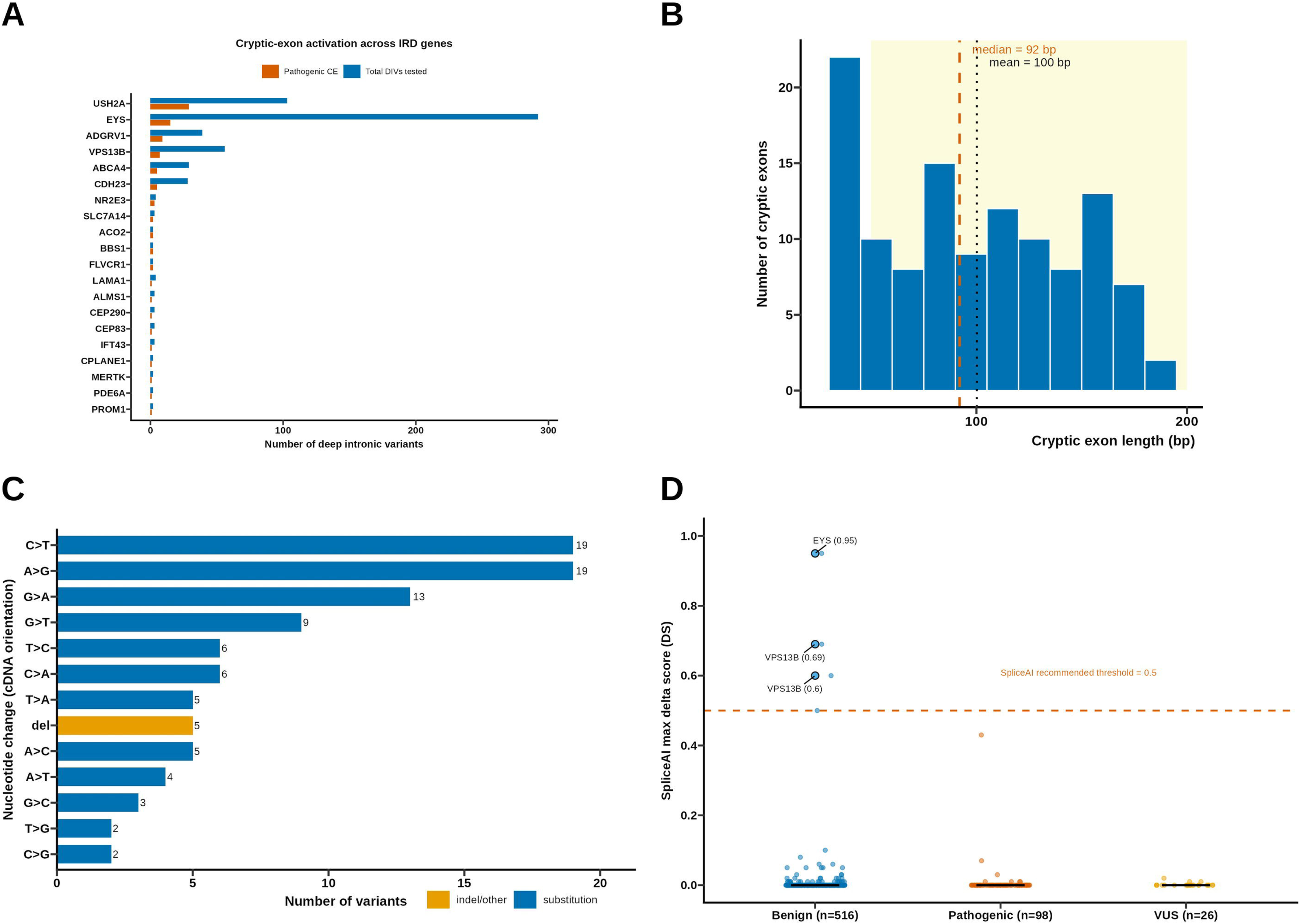
Cryptic-exon activation landscape among 640 deep intronic variants. (**A**) Deep intronic variants tested and pathogenic cryptic exons identified per gene (where >1 variants tested); USH2A and EYS contribute the largest burden. (**B**) Length distribution of activated cryptic exons (mean 100 ± 48 bp, median 92 bp; n = 117 measured CEs). (**C**) Spectrum of sequence changes among pathogenic variants (cDNA orientation); C>T and A>G transitions predominate. (**D**) SpliceAI maximum delta score by experimental class; 0 of 98 pathogenic variants reached the recommended 0.5 threshold, and the three high-scoring calls were experimental negatives.

The activated cryptic exons length ranged from 30 to 290 bp, with a mean of 100 ± 48 bp and a median of 92 bp (Figure 2B; n = 117 measured CEs). This distribution overlaps the 50–200 bp range most commonly reported for pathogenic pseudoexons in the literature,^53^ although the upper bound is constrained by the 270-bp insert and the true in vivo cryptic exon may extend beyond what the assay can display. In most cases a single dominant CE species accounted for >90% of aberrant reads, accompanied by minor alternative isoforms of different length, consistent with a principal cryptic splice-site choice plus lower-frequency alternatives. The staggered oligonucleotide design was important for this readout: for several variants the dominant cryptic exon was recovered only in the left-or right-shifted oligo, confirming that a single centered 270-bp window would have missed splicing events whose cryptic splice sites fall near an oligo boundary. This positional sensitivity also cautions that the assay yields a lower-bound estimate of activation, because a cryptic exon lying entirely outside all three windows would go undetected.

Among the pathogenic variants, single-nucleotide substitutions predominated over indels (Figure 2C). Collapsing complementary changes by strand, C>T/G>A (32) and A>G/T>C (25) transitions were the most frequent classes, followed by C>A/G>T (15), A>T/T>A (9), A>C/T>G (7), and C>G/G>C (5), with 5 deletions (Figure 2C). The predominance of C>T and A>G changes is consistent with the creation of new splice-donor motifs; a minority of variants generated splice-acceptor gains or acted through other mechanisms, including one intron-retention event (*ADGRV1* c.4379-97T>C). The six pathogenic deletions in *ADGRV1*, *CDH23*, *IMPG2*, *EYS*, *USH2A*, and *VPS13B* genes likely reshape local splicing regulatory context rather than directly introducing a canonical splice site, and their presence across six different genes indicates that small intronic deletions are a recurring, if minor, route to cryptic-exon activation that copy-number and single-nucleotide-variant pipelines are both poorly suited to flag. The overall bias toward donor-gain events is mechanistically informative: it suggests that, in this variant set, the rate-limiting step for pseudoexon inclusion is more often creation of a 5′ splice donor within otherwise permissive intronic sequence than creation of a 3′ acceptor, echoing the “wild-type pseudoexon” model in which latent acceptor-like elements await a newly minted downstream donor.^63^

### Current in silico predictors miss experimentally validated cryptic-exon variants

We asked whether the seven predictors in routine use could have identified the experimentally pathogenic variants. Focusing on SpliceAI, which performed best of the tools evaluated, we took the maximum delta score across the four SpliceAI predictions (acceptor/donor gain and loss) per variant. Among the 98 experimentally pathogenic variants, none reached the recommended pathogenic threshold of 0.5, only one reached 0.2 (*ABCA4* c.2160+998T>C, 0.43), and 91 of 98 (93%) scored exactly 0 (Figure 2D). Across the 26 VUS the maximum score was 0.02.

Strikingly, the only variants with high SpliceAI scores were experimental negatives. Three variants scored > 0.5 *EYS* c.6079-12630_6079-12627dup (0.95), *VPS13B* c.3446-11797T>G (0.69), and *VPS13B* c.148-17A>G (0.60) and all three failed to activate a cryptic exon in the assay, i.e., they are SpliceAI false positives (Figure 2D). The remaining six predictors: Pangolin, SQUIRLS, CI-SpliceAI, SPiP, MaxEntScan, and CADD performed worse than SpliceAI on this variant set; their scores were uninformative for deep intronic pathogenicity and are therefore not tabulated. Taken together, at standard operating thresholds current predictors would have discarded essentially all of the true cryptic-exon–activating variants recovered here, while their few confident calls were not reproduced experimentally.

### Orthogonal validation in extended genomic context and by segregation

Because the 270-bp assay context is short relative to native introns, we validated the classification in a longer, more physiological setting. A random subset of 10 pathogenic, 10 uncertain, and 10 benign variants was re-tested in a 1,000-bp genomic context and assayed by RT-PCR, with band intensities normalized to gel input to approximate the fraction of aberrant splicing (Figure 3). Eight of ten pathogenic variants (80%) were confirmed to activate a cryptic exon; the two that were not require further evaluation. Five of ten VUS (50%) produced visibly fainter aberrant bands, consistent with partial or context-dependent effects, and none of the ten benign variants (0%) activated a cryptic exon (Figure 3). The concordance at the extremes: high confirmation of pathogenic calls and complete absence of signal among benign calls supports both the assay and the chosen ALT/REF cut-offs, while the intermediate behavior of the VUS group underscores why 2–100-fold changes warrant additional testing rather than immediate classification.

**Figure 3.**
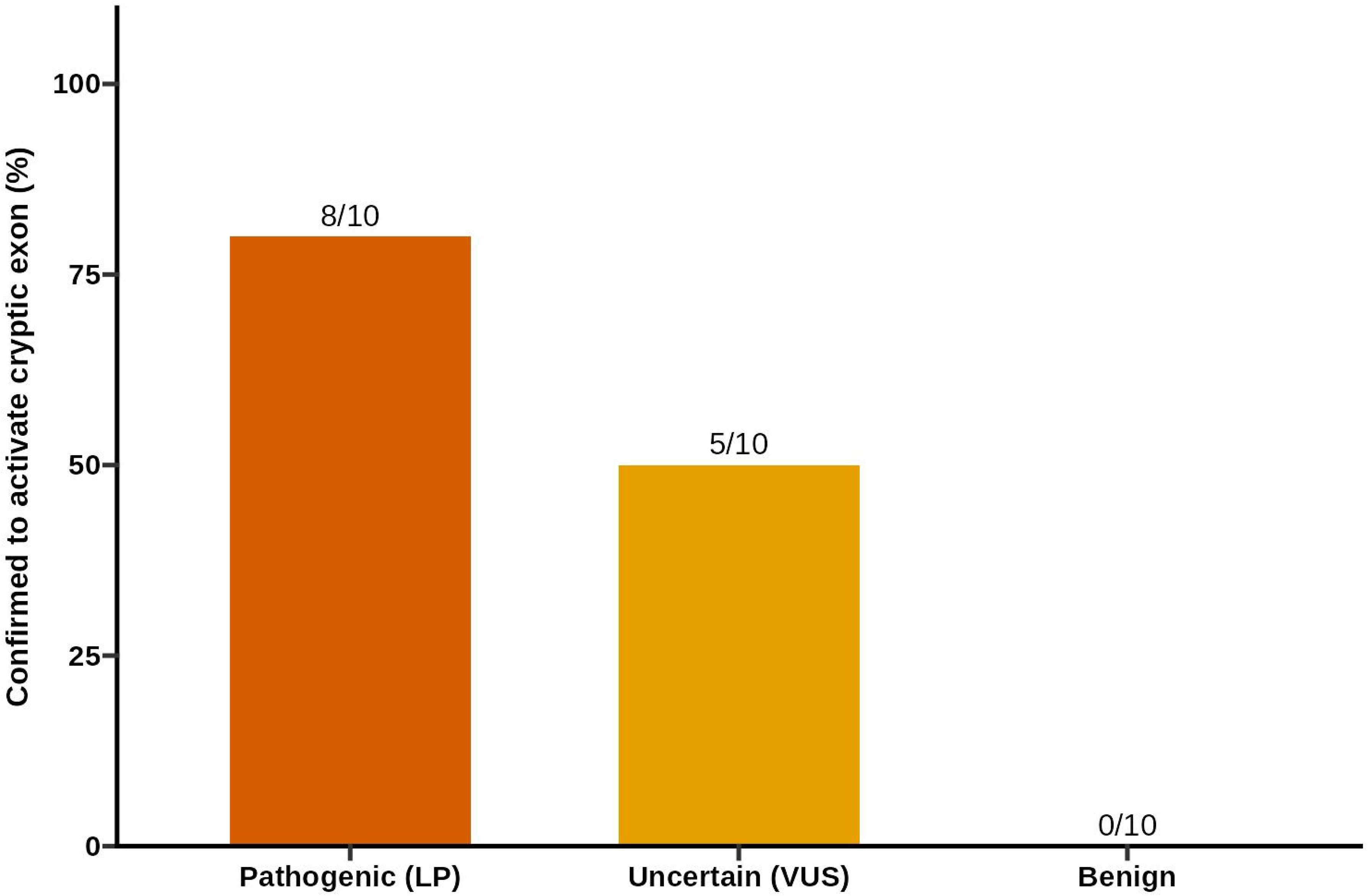
Orthogonal validation in a 1,000-bp genomic context. Confirmation rates by group for a random subset re-tested by RT-PCR: pathogenic 8/10 (80%), VUS 5/10 (50%), benign 0/10 (0%).

### Clinical impact: half of solved patients carry a single second-hit variant

Combining the functional and segregation evidence, cryptic-exon–activating variants provided a candidate second allele in 50 of the 78 patients (64%) analyzed. Among these 50 patients, the distribution of pathogenic DIVs was evenly split: 25 patients carried exactly one CE-activating variant, completing the biallelic diagnosis, whereas 25 patients carried more than one pathogenic DIV in the relevant gene. Multi-hit cases were most frequent in the long genes (*USH2A*, *EYS*, *ADGRV1*, *VPS13B*), where several rare DIVs may pass frequency filters and independently activate splicing. In these instances we prioritized the causal allele by integrating three lines of evidence: 1) concordance with the specific clinical phenotype, 2) family segregation establishing the *trans* configuration, and 3) the extended-context validation described above rather than assuming a single explanation from the screen alone.

## Discussion

Applying an unbiased high-throughput splicing assay to 640 patient-derived deep intronic variants, we found that 19.4% activated a cryptic exon, providing a candidate second allele for 64% of previously unsolved recessive IRD patients. Two findings have direct implications for how non-coding variants are interpreted in Mendelian disease. First, deep intronic pathogenicity is far more common among carefully selected “second-hit” candidates than prediction-based estimates would suggest.^64–66^ Second, the *in silico* tools currently used to triage these variants both miss the true positives and generate false positives, so empiric testing is necessary.

The predictor results deserve emphasis because they run counter to a common assumption. It has been argued that deep intronic variants contribute little to IRD causality,^7^ but that inference rests on computational filters that, in our data, assigned a score of exactly zero to 93% of experimentally pathogenic variants and never exceeded 0.5 for any of them. Conversely, the only variants SpliceAI scored confidently were experimental negatives. The apparent scarcity of pathogenic DIVs is therefore at least partly an artifact of the tools used to look for them: variants are filtered out before they can be tested, and their absence from curated datasets then reinforces the low scores, since tools such as SpliceAI are periodically retrained on accumulating labels.

Empiric datasets like this one is what is needed to recalibrate predictors,^45,67^ and until that recalibration matures, a near-zero SpliceAI score cannot be taken as evidence against pathogenicity for a deep intronic variant.

This has a concrete consequence for clinical variant curation. Under current ACMG/AMP guidelines, in silico evidence (PP3/BP4) is often applied using SpliceAI thresholds, and functional assays supply strong evidence (PS3/BS3). Our data indicate that for deep intronic positions the computational codes are effectively uninformative in both directions: a low score should not be read as benign supporting evidence (BP4), and the rare high score is not reliably corroborated by function, so the functional readout must carry the classification.^68–69^ The practical recommendation is that a rare deep intronic variant found *in trans* with a known pathogenic allele in a phenotype-matched recessive gene warrants direct splicing assessment regardless of its predictor scores, rather than being deprioritized because it scores zero. The false-positive calls are equally instructive: two of the three high-scoring variants were in *VPS13B* and *EYS*, genes with large introns in which SpliceAI evidently identifies motif-level donor or acceptor gains that are not realized as exon inclusion in a cellular context, a reminder that a predicted splice-site change and an actual splicing outcome are not the same measurement.

The scale of the benign fraction is itself a useful resource. The 516 variants that did not alter splicing constitute a large, uniformly assayed set of experimentally neutral deep intronic changes in disease-relevant genes - the kind of negative data that supervised splicing models are typically insufficient. Sharing both the positive and negative classifications, as we do here, should improve the calibration of future tools more than positive examples alone. However, our HTSA experimental method was sharpened to detect cryptic exon activation, so it might miss some other effects that these intronic variants might cause.^70^ Further investigation is needed.

Gene architecture shapes both the yield and the difficulty of this approach. Seventy-one percent of pathogenic variants fell in the six longest genes, and multi-hit cases clustered there as well. In short genes with few rare DIVs variant choice is straightforward and an extremely rare DIV that passes filtering is often the answer. *IMPG2* is a clear example of that. In long genes such as *USH2A*, *EYS*, *ADGRV1*, and *VPS13B*, by contrast, multiple rare DIVs may each activate splicing in the assay, and distinguishing the causal allele requires phenotype concordance, segregation, and additional validation. Consistent with prior mapping of cryptic-exon–prone regions in *USH2A*,^19,21^ the pathogenic DIVs we recovered were not evenly distributed along the introns, suggesting that specific intronic subregions are especially susceptible and could be prioritized in future, denser screens.

Several limitations follow from the assay design. The 270-bp insert constrains the observable cryptic-exon length and its local context, so a CE detected here may be longer, shorter, or spliced differently in the native locus; the extended 1,000-bp validation mitigates but does not eliminate this concern, and the two pathogenic variants not confirmed at 1,000 bp illustrate the point.

Transient transfection produces a high plasmid copy number and some background splicing, which is why we relied on the ALT/REF ratio rather than absolute ALT signal. We also evaluated a stable, chromosomally integrated landing-pad format with FACS-based sorting of GFP-positive and GFP-negative cells; although attractive in principle for single-cell quantification, we found the false-positive rate during sorting unacceptably high and did not adopt it. Finally, the assay reports on splicing in HEK293T cells and cannot capture retina-specific splicing factors^71^; the degree of aberrant splicing, and therefore the clinical severity, may differ in the target tissue, and the phenotypic spectrum can plausibly track the fraction of transcripts that are mis-spliced.

These caveats notwithstanding, the therapeutic relevance is direct. Pseudoexon-activating variants are among the most tractable targets for splice-modulating antisense oligonucleotides (ASOs), which can block the aberrant splice site and restore normal exon usage, as demonstrated for *ABCA4, CEP290, DMD*, *USH2A* and some other genes.^11,16,72–73^ Because an ASO must be designed against the specific cryptic splice junction, a molecular diagnosis resolved to the level of the individual cryptic exon is the prerequisite for a candidate intervention.^74–75^ The assay reported here delivers that granularity: for each pathogenic variant it returns the sequence and boundaries of the activated exon, which can be read directly into ASO or gene-editing design.

This is especially consequential for the long genes that dominated our pathogenic set (*USH2A*, *EYS*, *ADGRV1*), which exceed the cargo capacity of adeno-associated virus and are therefore poor candidates for gene augmentation but are well suited to splice correction.

The clinical workflow implied by these findings is a staged one. WGS identifies unsolved recessive cases with a single P/LP allele; rare deep intronic variants in the matching gene are enumerated without regard to predictor scores; the HTSA classifies their splicing effect; and phenotype concordance plus segregation establish the *trans* configuration and resolve multi-hit genes. The even split we observed half of solved patients with a single CE-activating variant and half with several, shows that both the simple and the complex scenarios are common, and that a decision process combining functional, genetic, and clinical evidence is needed rather than a rule that assumes one variant per case. As WGS becomes routine for unsolved recessive disease, we anticipate that empiric splicing assays will be paired with prediction and standardized in diagnostic pipelines, supported by robust analysis that classify DIVs at scale^76^ and by reference sets of experimentally validated pathogenic and benign variants. Although developed for IRDs, nothing in the approach is retina-specific: the same logic applies wherever a recessive Mendelian disorder is left with a single coding allele and an intron full of uninterpreted rare variants, and the recurrent under-recognition of deep intronic pathogenicity across genetic medicine^17–18,22^suggests the yield elsewhere could be comparably high. The data presented here are intended to support that transition and to improve the interpretation of non-coding variation in the retinal disease genes and beyond.

## Conclusions

Use of an unbiased approach to assess the pathogenicity of as many intronic variants as possible to accurately identify novel CEs is crucial. Overall, 124 of the 640 variants (19.4%) tested activated CEs. We hope these empiric data regarding variant pathogenicity and additional data can be used to improve interpretation of sequence variants identified in patients, and to improve algorithms for assessment of non-coding variant pathogenicity.

## Data Availability

Raw data are available for re-analysis on reasonable written request to the corresponding author (M.W.).

## Acknowledgments

The authors thank all subjects for their participation in this study and the OGI Genomics Core members for their experimental assistance. The authors also acknowledge Jason Comander for valuable input and advice on data analysis, future experimental design, and research directions.

## Author Contributions

M.W., K.B., and E.A.P. conceptualized and planned the experiments. M.W. executed experiments and wrote the manuscript; K.B. and E.A.P. reviewed and edited. S.M. and D.N. carried out bioinformatics analysis. K.V. performed part of the experiments. H.S. provided NGS services and experimental protocol improvements. M.W. and K.B. carried out the formal analysis. E.M.P. curated data from IRD patients by performing genetic counseling. All authors provided critical feedback on the manuscript.

## Ethics Declaration

The study was approved by the Institutional Review Board at Massachusetts Eye and Ear (Human Studies Committee MEE, Mass General Brigham, USA) and adhered to the tenets of the Declaration of Helsinki. Informed consent was obtained from all individuals on whom genetic testing and further molecular evaluations were performed.

## Abbreviations

AD: autosomal dominant
AR: autosomal recessive
CE: cryptic exon
DIV: deep intronic variant
HTSA: high-throughput splicing assay
IRD: inherited retinal disorder
NGS: next-generation sequencing
WES: whole-exome sequencing
WGS: whole-genome sequencing

---

**Supplementary Table S1.** Oligonucleotides (REF and ALT) for the 640 variants tested in this study (672 total, 32 excluded from final analysis), with read counts for three biological replicates and corresponding SpliceAI prediction scores.

